# Inferring inter-chromosomal rearrangements and ancestral linkage groups from synteny

**DOI:** 10.1101/2023.09.17.558111

**Authors:** Alexander Mackintosh, Pablo Manuel Gonzalez de la Rosa, Simon H. Martin, Konrad Lohse, Dominik R. Laetsch

## Abstract

Chromosome rearrangements shape the structure of the genome and influence evolutionary processes. Inferring ancestral chromosomes and rearrangements across a phylogenetic tree is therefore an important analysis within evolutionary genetics. One approach to this inference problem is to focus on synteny information, i.e. the co-occurrence of loci on the same chromosome. Although algorithms for inferring ancestral linkage groups (ALGs) and inter-chromosomal rearrangements from synteny have been previously described, they have seldom been applied to modern genome data. Here we implement these algorithms in a command-line tool, `syngraph`, and evaluate their performance using simulations that include a mix of different rearrangements and types of error. We show that ALGs and rearrangements can be recovered when the rearrangement frequency per-branch is well below the number of chromosomes. We demonstrate that competing models of rearrangement can be inferred by comparing observed results to simulations. Finally, we reanalyse genome assemblies of rhabditid nematodes and find that independent fusions of the same ALGs pose a challenge that is difficult to overcome without gene-order information. Our simulations and analysis of real data demonstrate both the promise and limitations of using synteny information to infer patterns of genome evolution.

## Introduction

The fact that the genomes of different organisms vary in chromosome number and structure has long been appreciated (Robertson 1916; Sturtevant 1921). Changes in ploidy explain at least some of this variation, especially in plants (Otto and Whitton 2000). Chromosome rearrangements are another mechanism by which genomes undergo large-scale changes, and they are common across eukaryotes (Zhao and Schranz 2019; Li *et al*. 2022; Muffato *et al*. 2023). Intra-chromosomal rearrangements involve a single chromosome (e.g. inversions) while inter-chromosomal involve two chromosomes (e.g. fissions, fusions, and translocations). Chromosome rearrangements can influence fundamental evolutionary processes, such as recombination (Bidau *et al*. 2001; Näsvall *et al*. 2023), as well as broader processes like speciation (Yoshida *et al*. 2023; Mackintosh *et al*. 2023), so there is considerable interest in reconstructing how genomes have rearranged through time. Ancestral karyotypes and rearrangements have been estimated for a number of different taxa, including *Drosophila*, ruminants, and birds (Muller 1940; Farré *et al*. 2019; Damas *et al*. 2018), as well as large taxonomic groups such as rosid plants and animals (Murat *et al*. 2015; Simakov *et al*. 2022). Typically, such analyses rely on genome sequences or linkage map data for tens of genomes (or less), but it is becoming more common to analyse hundreds of chromosome-level genome assemblies (Muffato *et al*. 2023; Wright *et al*. 2023), highlighting the need for efficient inference methods.

Estimating ancestral genomes and rearrangements given a set of present-day genomes and a phylogenetic tree is a challenging combinatorics problem (Sankoff 2003; Fertin *et al*. 2009). Methods for inferring ancestral genomes are typically either event-based or adjacency-based (Feng *et al*. 2017). Event-based methods use an explicit model of rearrangement and aim to construct maximally parsimonious ancestral genomes at internal nodes by minimising the number of rearrangements across the tree (e.g. Bourque and Pevzner 2002; Zheng and Sankoff 2011). By contrast, adjacency-based methods reconstruct ancestral genomes by assuming that genome structure that is conserved in present-day genomes was also present in their most recent common ancestor, without explicitly modelling individual rearrangements (e.g. Kim *et al*. 2017; Muffato *et al*. 2023). The event-based approach has the advantage of co-estimating ancestral genomes and rearrangements, and so gives direct insight into the evolutionary process, but comes at a computational cost. These approaches can therefore be viewed as complementary, with adjacency-based analyses being the most practical way to summarise patterns of genome evolution from hundreds of genomes (Muffato *et al*. 2023) and event-based approaches being a better choice for detailed rearrangement inference from a handful of genomes (Ostevik *et al*. 2020).

Genome evolution through time can be reconstructed at different resolutions; the most detailed being the full reconstruction of ancestral sequences. However, co-estimating base and indel substitutions and chromosome rearrangements requires accurate whole genome alignments and therefore considerable computational resources (Armstrong *et al*. 2020). An alternative is to instead infer the order of sequences in ancestral genomes without considering substitutions. This task is straightforward when gene-order is well conserved, but becomes more challenging when the intra-chromosomal rearrangement rate is high or if taxa are very distantly related (Farré *et al*. 2019; Muffato *et al*. 2023). Genome evolution can also be reconstructed at the level of synteny (Fertin *et al*. 2009). While the term synteny is sometimes used to denote co-linearity between chromosomes, here we use it to refer to the co-occurrence of loci on the same chromosome, regardless of their order (Renwick 1971; Passarge *et al*. 1999). Focusing exclusively on synteny means that only unordered sets of markers (i.e. linkage groups) and inter-chromosomal rearrangements can be reconstructed. Despite this limitation, methods that focus on synteny are likely to be applicable across a wide range of datasets, as synteny decays more slowly than gene-order across evolutionary time (Simakov *et al*. 2022). An efficient synteny-based method for reconstructing genome evolution would therefore allow for the processes that constrain and promote fission, fusion and translocation rearrangements to be investigated across many groups of species.

Although algorithms exist for reconstructing genome evolution from synteny (Ferretti *et al*. 1996; DasGupta *et al*. 1997; Liben-Nowell 2001), they have rarely been applied to data. Additionally, it is not clear how well these methods perform under different rearrangement scenarios or how well they can accommodate the types of error that exist in genome assemblies and annotations. Here we address these issues by implementing previously described algorithms in a command-line tool – `syngraph` – and performing analyses on both simulated and real data. The manuscript is structured as follows: First, we briefly recapitulate previously described synteny-based algorithms for inferring inter-chromosomal rearrangements between genomes and across phylogenies. Next, we evaluate the performance of these methods on data simulated under a range of rearrangement and error parameters. Finally, we reanalyse a set of nematode genomes from Gonzalez de la Rosa *et al*. (2021) and compare our results to theirs.

## Results

### Inferring inter-chromosomal rearrangements between two genomes

Ferretti *et al*. (1996) outlined the problem of calculating a syntenic edit distance between two genomes (hereafter simply referred to as syntenic distance). This is the minimum number of fissions, fusions and translocations required to transform one genome into another (Figure 1). The entire genome sequences are not required, instead markers present in both genomes only need to be assigned to chromosomes in each and positional information (i.e. marker order) is ignored. A marker can be any single-copy sequence feature that is identifiable across both genomes, e.g. ultra-conserved elements, genes or nucleotide alignments. Given this information, Ferretti *et al*. (1996) showed that the problem of transforming genome *G*_*A*_ to genome *G*_*B*_ by rearrangement can be reduced as follows: write each chromosomes of *G*_*A*_ as a set populated by the labels of chromosomes in *G*_*B*_ with which it shares markers. Then, *G*_*A*_ must be rearranged through the smallest number of set operations such that the final sets are all unique and of length one (therefore each representing a single chromosome from *G*_*B*_). In this scheme, fusions are unions of two sets and fissions replace one set with two disjoint sets (Ferretti *et al*. 1996). Additionally, a translocation can be modelled as an exchange of subsets (Ferretti *et al*. 1996). In principle, these set operations can be combined to find a maximally parsimonious series of rearrangements. Calculating the syntenic distance between two genomes therefore gives (i) a measure of how rearranged they are from one another and (ii) a putative history of the rearrangements between them (Figure 1).

**Figure 1.**
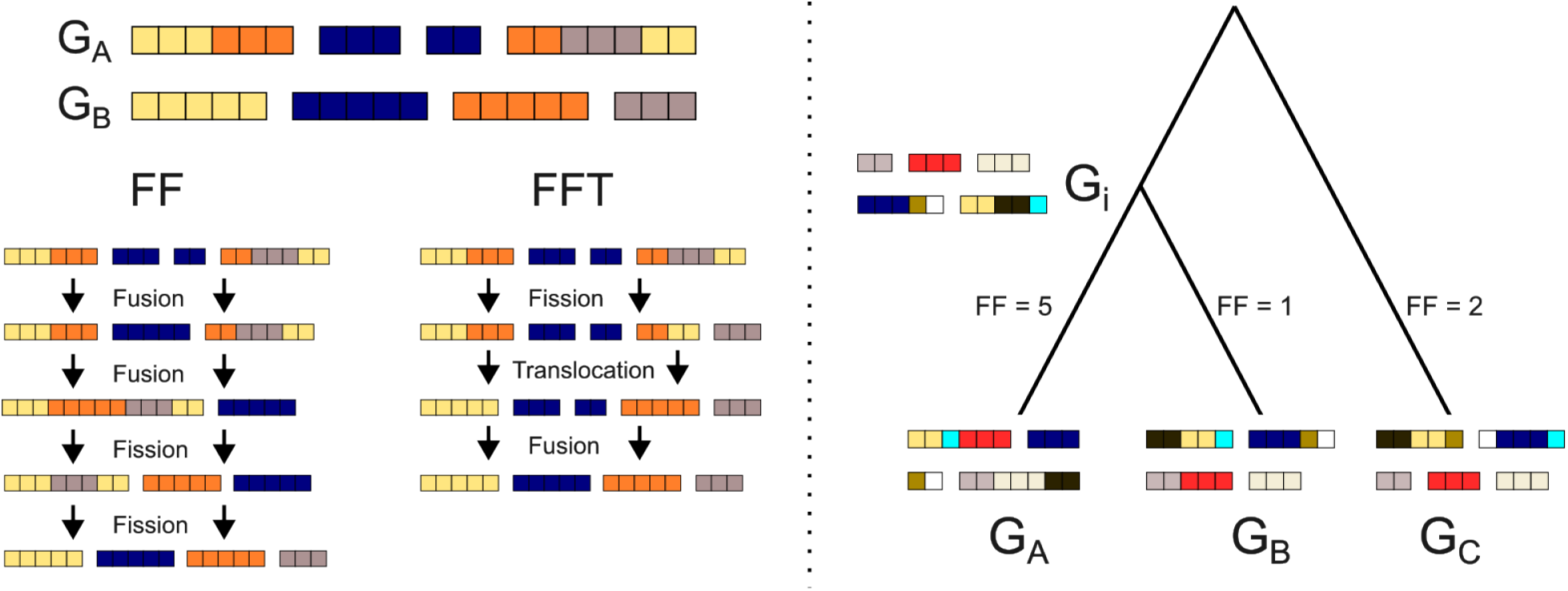
Inferring rearrangements between two genomes and across a phylogenetic tree. The left panel shows two genomes, *G*_*A*_ and *G*_*B*_, where chromosomes (rectangles) consist of markers (squares), and the order of markers within a chromosome is arbitrary. Each marker in *G*_*A*_ is coloured by the chromosome that it is found on in *G*_*B*_. Given this representation of *G*_*A*_, the FF and FFT algorithms can be used to transform it to *G*_*B*_, thus calculating a syntenic distance between *G*_*A*_ and *G*_*B*_. The right panel shows three genomes, *G*_*A*_, *G*_*B*_ and *G*_*C*_, that are related by a phylogenetic tree. Here markers are coloured by the synteny-set that they are a part of, where a synteny-set is defined as a group of markers that are syntenic in genomes *G*_*A*_, *G*_*B*_ and *G*_*C*_. A genome at the internal node of the tree, *G*_*i*_, can be constructed by combining synteny-sets and evaluated by summing the syntenic distances between *G*_*i*_ and the other three genomes. Branches of the phylogeny are labelled with the syntenic distance under the FF algorithm given the genome at *G*_*i*_.

We implemented two heuristic algorithms for calculating a syntenic distance between two genomes: the algorithm from DasGupta *et al*. (1997) where only fission and fusion rearrangements are permitted (FF) and a modified version of the algorithm from Ferretti *et al*. (1996) which allows fission, fusion and translocation (FFT) (Figure 1). While Ferretti *et al*. (1996) include the possibility of non-reciprocal translocations in their algorithm, we only consider reciprocal translocations where portions of both chromosomes are exchanged. The FF algorithm is straightforward: chromosomes of *G*_*A*_ that share markers syntenic in *G*_*B*_ are fused recursively, then fissions are implemented to recover the individual chromosomes of *G*_*B*_ (Figure 1). The FFT algorithm uses similar criteria for implementing fissions and fusions, but will also implement a translocation when two chromosomes share multiple sets of markers syntenic in *G*_*B*_ (Figure 1). Given that this algorithm is more complex, we do not describe the details here and instead provide a description in the Supplementary Methods.

We used simulations to test whether the syntenic distances estimated by these algorithms correspond to the true number of total rearrangements. More specifically, we simulated an initial genome containing 1,000 markers uniformly distributed among *k* chromosomes and then generated a second genome by rearranging the first *r* times. Note that here we only simulated the types of rearrangement considered by the respective algorithms. We performed simulations for three values of *k* (10, 20 and 30) and varied *r* between 1 and 50. We find that both algorithms accurately estimate the total number of rearrangements when *r <*= *k* (Figure 2). Above this, both algorithms infer syntenic distances that are underestimates of the total number of rearrangements (Figure 2). This bias is much more pronounced when simulations and inference include translocation, although this is expected given that a series of fissions and fusions can sometimes be explained by a single translocation. Put differently, histories that contain all three types of rearrangements are more challenging to estimate. Nonetheless, these results suggest that the syntenic distance between two genomes is a useful approximation of the true number of total rearrangements, as long as the total number of rearrangements is less than the number of chromosomes.

**Figure 2.**
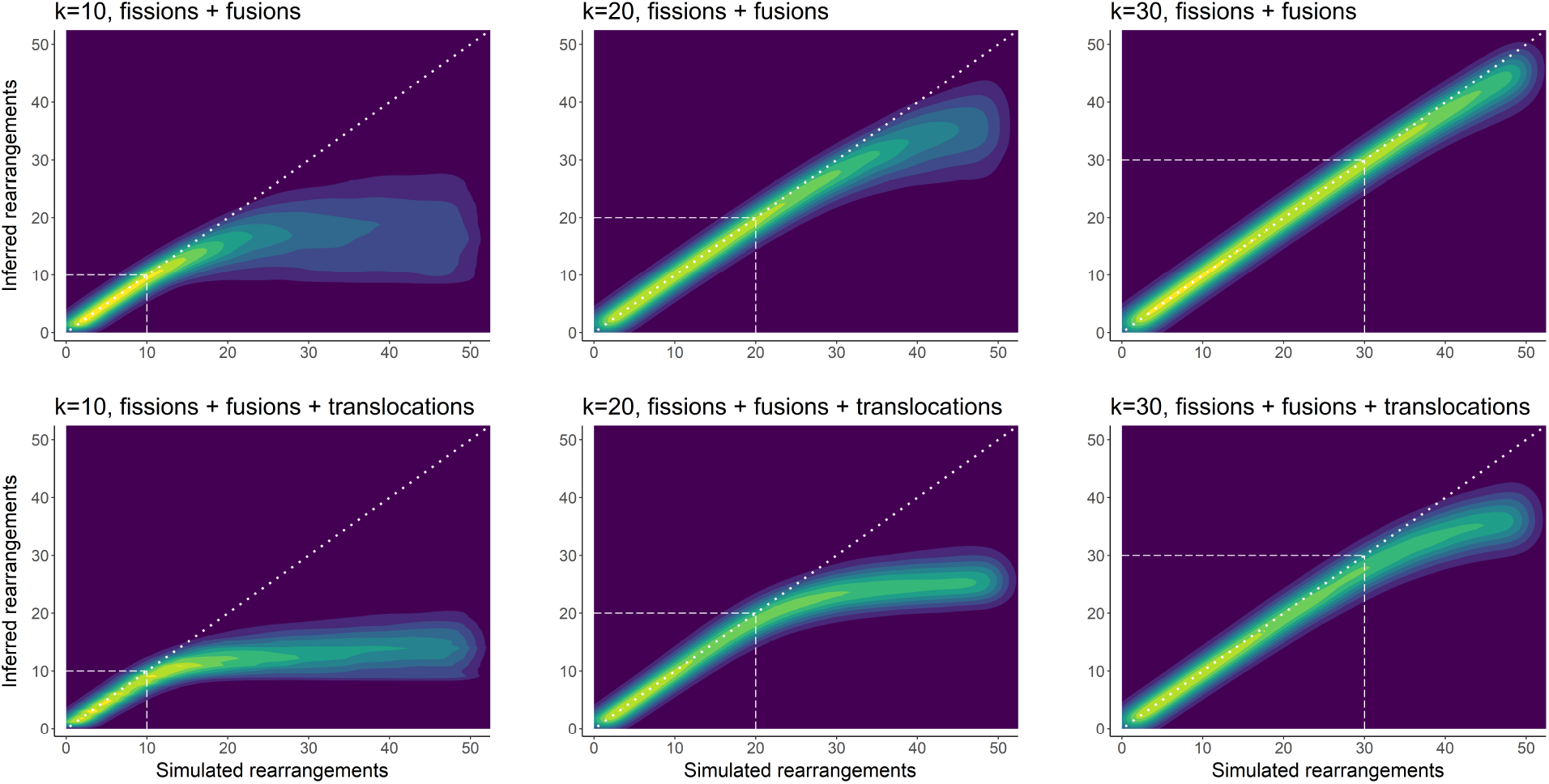
Estimating the total number of rearrangements between two genomes using syntenic distance. Each plot summarises results from 50,000 simulations, with brighter colour corresponding to a greater density of simulated data points. The total number of simulated rearrangements is plotted on the x-axis with the inferred number on the y-axis. The dotted white line along the diagonal (*x* = *y*) corresponds to inferred and simulated counts being equal, i.e. correct inference. Plots show simulations with *k* = 10 (left), *k* = 20 (middle), or *k* = 30 chromosomes (right). Additionally, plots either show simulations containing only fissions and fusions (top) or fissions, fusions and translocations (bottom). Dashed lines in each plot show where the number of rearrangements is equal to *k*.

### Inferring ALGs and inter-chromosomal rearrangements across a phylogeny

The syntenic distance between genomes can be used to reconstruct ancestral linkage groups (ALGs) and inter-chromosomal rearrangements across a phylogenetic tree (Ferretti *et al*. 1996; DasGupta *et al*. 1997). Consider a triplet of related genomes, *G*_*A*_, *G*_*B*_ and *G*_*C*_, where *G*_*A*_ consists of chromosomes *A*_1_, *A*_2_, …, *A*_*n*_ with arbitrary labels. Any marker present in all three genomes can be written as the chromosomes it is found on, e.g. [*A*_4_, *B*_22_, *C*_9_]. Markers that are syntenic in all three genomes will have the same notation as each other and can be considered as part of the same synteny-set (Figure 1). These synteny-sets can be used as building blocks to reconstruct the ALGs of *G*_*i*_ (the ancestral genome at the internal node of the tree relating *G*_*A*_, *G*_*B*_ and *G*_*C*_) (Figure 1). An adjacency-based approach to ALG reconstruction is to initiate a LG with a single synteny-set and add all synteny-sets that are syntenic with it in two of the triplet genomes (i.e. those that share two of the three elements in their notation), and repeat until all synteny-sets are part of a LG. An event-based approach is to build many different versions of ALGs from synteny-sets under more relaxed rules (e.g. synteny-sets in the same LG are allowed to share just one element in their notation) and then identify the most parsimonious set of ALGs using the sum of syntenic distances between *G*_*i*_ and *G*_*A*_, *G*_*B*_, *G*_*C*_. Once a set of ALGs is obtained for *G*_*i*_ (by either method), rearrangements can be recorded between *G*_*i*_ and its descendants (*G*_*A*_ and *G*_*B*_) using the FF or FFT algorithms.

To assess these approaches to ALG estimation, we performed simulations over a phylogenetic tree with *n* = 3 leaves (see Methods for details). The genome at the root of the tree always had 20 chromosomes and 1000 markers (these parameters are used for all subsequent simulations), and the number of rearrangements simulated across the tree varied between 1 and 50. Given the synteny of markers in the three sampled genomes, we estimated ALGs and rearrangements with the approaches described above and calculated two performance metrics:

- **ALG accuracy**: The proportion of markers within correctly estimated ALGs, averaged across all internal nodes of the phylogeny (unless stated otherwise).
- **Rearrangement accuracy**: The proportion of branches across the tree with the correct number of estimated rearrangements by type, e.g. 1 fission, 2 fusions, 1 translocation.

We find that both ALG and rearrangement accuracy are greater when simulations and inference only contain fissions and fusions (Figure 3). Both metrics decline with the number of simulated rearrangements (Figure 3), but rearrangement accuracy does so faster (note the difference in y-axes within Figure 3). This is expected given that poor rearrangement estimation will only typically involve a subset of ALGs, meaning that some ALGs (e.g. those that are invariant across the tree) can still be well estimated when rearrangements are not. These simulations also show that rearrangements across the tree are only well estimated when their frequency per-branch is well below the number of chromosomes (in this case *∼* 20, Figure 3). Surprisingly, we find that the event-based approach often gives worse results than the simpler and quicker adjacency-based approach (Figure 3). We therefore use adjacency-based ancestral genomes for all subsequent analyses.

**Figure 3.**
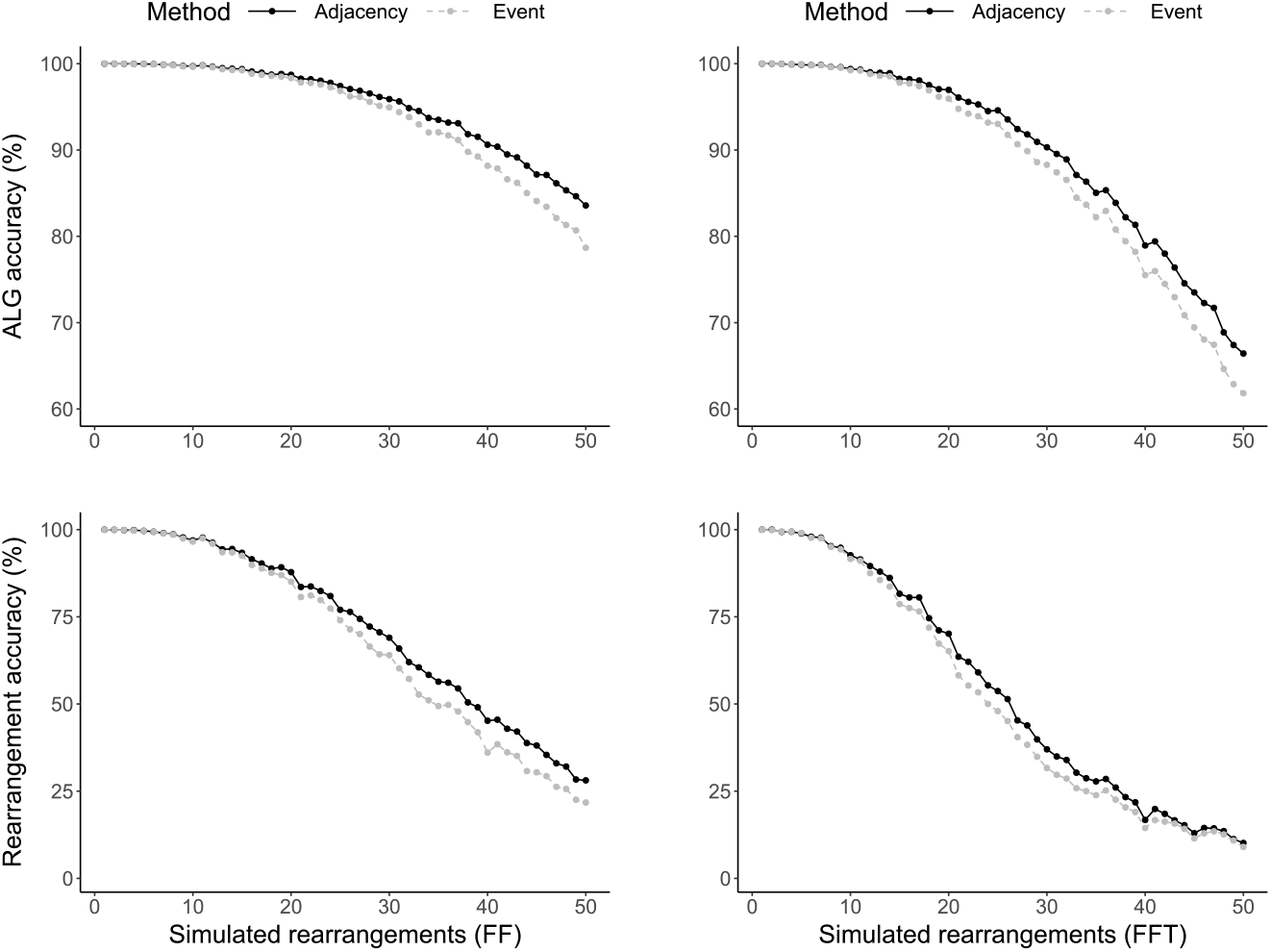
The accuracy of inferred ALGs and rearrangements across a tree with *n* = 3 leaves. The top plots show the accuracy of ALGs inferred from simulations with between 1 and 50 rearrangements. The bottom plots show the accuracy of inferred rearrangements. Points always represent averages from 1000 simulations. Simulations either include fissions and fusions (left) or fissions, fusions, and translocations (right). Whether ALG estimation was performed using an adjacency or event-based approach is shown using colour and line-type. Note the difference in y-axis between the plots showing ALG accuracy (60-100%) and rearrangements accuracy (0-100%).

The approach outlined for a phylogeny with *n* = 3 leaves can be extended to arbitrarily large trees. To do this, nodes are visited through a post-order traversal (from leaves to root) and ancestral genomes are reconstructed using a triplet of genomes (either observed genomes at the leaves or already inferred ancestral genomes). This process continues until the ancestral genomes at all internal nodes have been estimated. Given a large phylogeny (*n* = 100 leaves), we tested how well ALGs are estimated at a deep node in the tree (the child-node of the root with the most descendants). When the total number of fission and fusions rearrangements simulated was 198, 594 and 990, corresponding to an average of 1, 3 and 5 rearrangements on each of the 2*n −* 2 branches, ALG accuracy (under the FF algorithm) was 98.53%, 67.48%, and 13.94%, respectively. These performance estimates are far worse than analogous ones for a tree with only *n* = 3 leaves (99.98%, 99.65% and 98.72% for 1, 3 and 5 rearrangements per-branch, respectively). This shows that errors in ALG estimation accumulate upwards through the tree, especially when the rearrangement rate is high.

### The effect of marker error

We have so far only evaluated the performance of ALG and rearrangement inference using simulations that produce perfect data. Real genomes sequences and annotations, however, are likely to contain errors. For example, some markers will be missing from certain genomes and some markers may be assigned incorrect orthology. We therefore added these sources of error to simulations.

We implemented a method to assign markers that are missing in a minority of genomes to ALGs (see Methods). We then tested this method by modifying our simulations so that genomes at the leaves of the tree have a small number of missing markers. We simulated fissions and fusions over a tree with *n* = 10 leaves and inferred them back under the FF algorithm. We find that, even when the amount of missingness is small (e.g. 5% of markers per-genome), ALG accuracy is significantly reduced (Figure 4). By contrast, missingness only has a small effect on whether rearrangement histories are estimated accurately (Figure 4). This disparity can be explained by the fact that missingness removes information about individual markers from the data, which effects our ability to estimate all of the markers within an ALG correctly but not our ability to identify whether, for example, a single fusion rearrangement has happened on a particular branch. Given the sensitivity of our ALG accuracy measure to missingness, we considered an alternative metric:

**Figure 4.**
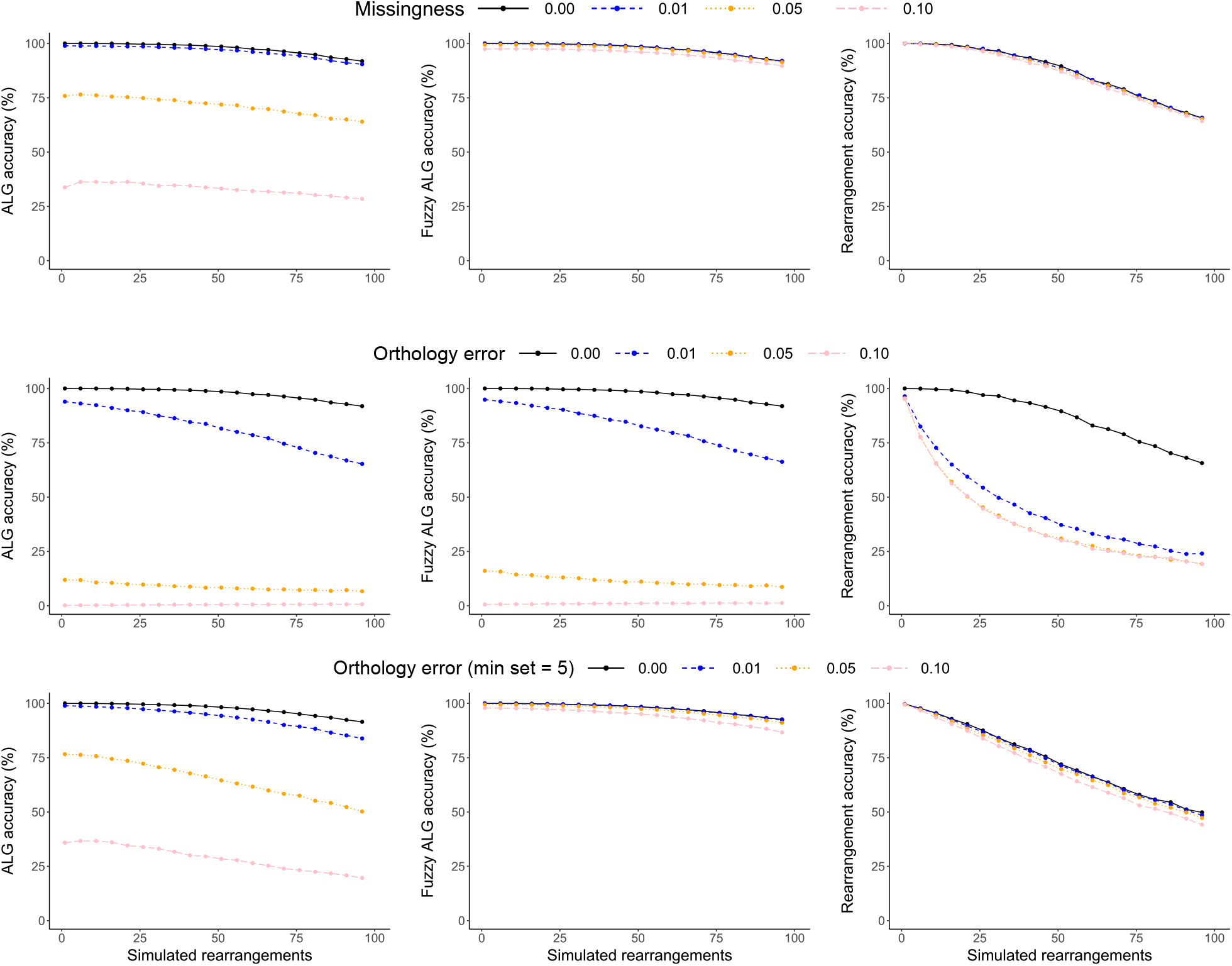
The effect of marker error and rearrangement frequency on inference accuracy. Each plot shows how the number of rearrangements (x-axis) affects a performance metric (y-axis). These metrics include ALG accuracy (left column), fuzzy ALG accuracy (middle column), and rearrangement accuracy (right column). The top row of plots include simulations with varying levels of marker missingness. The middle row of plots include different levels of marker orthology error, whereas the bottom row include the same levels of orthology error but inference was performed with a minimum synteny-set size of 5. Points always represent averages from 1000 simulations.

- **Fuzzy ALG accuracy**: The proportion of markers within fuzzily estimated ALGs, averaged across all internal nodes of the phylogeny (unless stated otherwise). A fuzzy ALG contains at least 90% of markers from a single true ALG and no more than 5% of markers from any other.

We find that fuzzy ALG accuracy is generally high at all levels of missingness (Figure 4). Rearrangements and good approximations of ALGs can therefore still be estimated despite missingness. We next added orthology error to simulations by randomising the chromosome assignment of a small proportion of markers. We again simulated rearrangements across a tree with *n* = 10 leaves and find that, perhaps unsurprisingly, orthology error has a large effect on all performance metrics (Figure 4). In particular, it becomes difficult to estimate rearrangement histories as erroneous rearrangements are introduced to account for the movement of markers (Figure 4). These inferred rearrangements will involve a much smaller number of markers than fission, fusion, or translocation events which typically involve large portions of chromosomes. We therefore set a minimum synteny-set size for inferring ALGs and rearrangements in an attempt to improve performance (see Methods). Setting the minimum synteny-set size to five markers resulted in large improvements in fuzzy ALG and rearrangement accuracy (Figure 4). However, even without error, the addition of a minimum synteny-set size reduces the frequency at which rearrangement histories are perfectly estimated (Figure 4). This is presumably due to an inability to identify fissions that involve a small number of markers, and a similar difficulty in reconstructing complex rearrangement sequences that result in small sets of syntenic markers. These simulations show that a small amount of orthology error can be overcome by introducing a minimum set size, although this itself does have a performance cost.

### Evaluating evidence for translocations through simulation

Only fission and fusions rearrangements are inferred when using the FF algorithm, even if the true rearrangement history contains translocations. The FFT algorithm, by contrast, allows inference of all three types of rearrangement, but it is not clear how often it erroneously infers a series of fissions and fusions as a translocation, or vice versa. We therefore investigated whether inference under the FFT algorithm recovers the correct ratio of fission, fusion and translocation events. We again simulated rearrangements over a tree with *n* = 10 leaves. We varied the number of rearrangements as well as the ratio of fission, fusion and translocation events (1:1:0, 1:1:1 or 1:1:2), then performed inference under the FFT algorithm. When only fission and fusion rearrangements are simulated (ratio 1:1:0), a small number of translocations are still inferred (Figure 5). For example, when 50 fissions / fusions are simulated across the tree, an average of 4.1% of inferred rearrangements are translocations, with 95% confidence intervals (95% CIs) of 0.0 and 12.2%. This shows that although the false-positive rate is generally low, rearrangement histories that include only fissions and fusions can result in inferred histories where *∼* 10% of rearrangements are translocations. When translocations are included in simulations with ratio 1:1:1 or 1:1:2, they are inferred at the expected frequencies of 33.3% and 50.0%, albeit with a downwards bias that increases with the number of simulated rearrangements (Figure 5). Although inferred rearrangements across many simulations do tend to reflect the underlying rearrangement ratio, the wide variation among simulation replicates (Figure 5) shows that a single analysis only contains limited information about the relative rates of fission, fusion and translocation.

**Figure 5.**
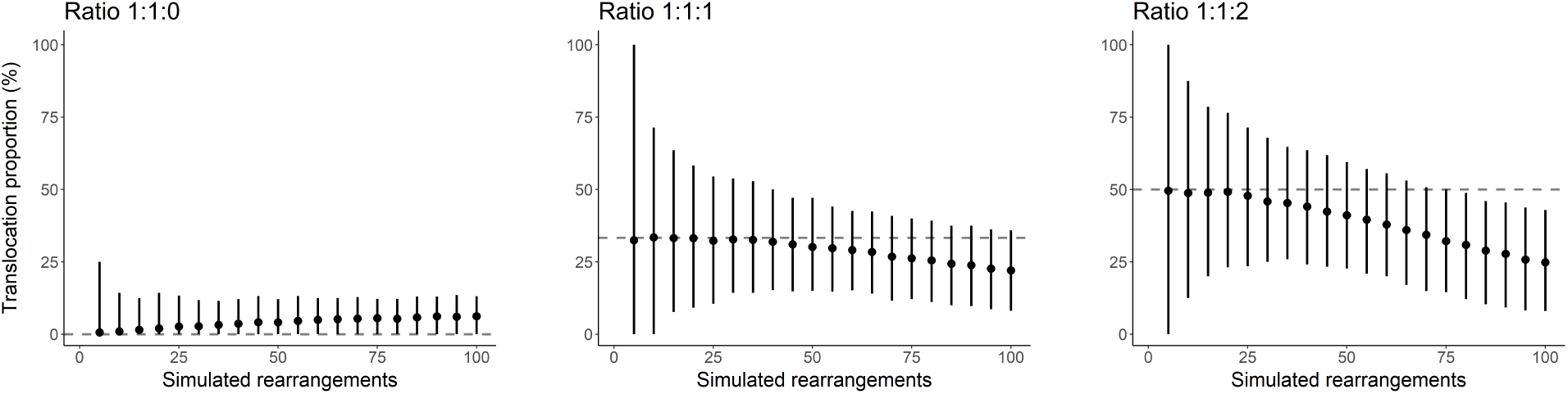
The proportion of rearrangements inferred as translocations for simulations across a tree with *n* = 10 leaves. Each plot shows the mean translocation proportion and 95% CIs (y-axis) for simulations involving different numbers of rearrangement (x-axis). The fission:fusion:translocation ratios of simulated rearrangements are 1:1:0 (left), 1:1:1 (middle), and 1:1:2 (right). Dashed horizontal lines correspond to the expected translocation proportion under each rearrangement ratio.

### Rhabditid nematodes and Nigon elements

We implemented the methods described above for inferring ALGs and rearrangements in a command line tool, `syngraph`. Here we apply `syngraph` to genomes of nematodes in the order Rhabditida. Species in this order typically possess a small number of chromosomes (*∼* 6), albeit with some exceptions (Table S1). Pairwise comparisons between genomes have shown that gene-order is highly variable across species (Lee *et al*. 2003; Hillier *et al*. 2007; Stevens *et al*. 2020). By contrast, synteny is more conserved between genomes and has therefore been used to identify seven ancestral linkage groups (ALGs), often referred to as Nigon elements, as well as inter-chromosomal rearrangements (Tandonnet *et al*. 2019; Gonzalez de la Rosa *et al*. 2021). We chose to reanalyse 14 rhabditid nematode genomes from Gonzalez de la Rosa *et al*. (2021). They used a clustering algorithm to identify ALGs and then manually inferred fissions and fusions across the tree (Gonzalez de la Rosa *et al*. 2021). By contrast, the synteny-based method we focus on aims to co-infer ALGs and rearrangements across a phylogeny in a single automated analysis.

We used BUSCO genes as orthologous markers for measuring synteny. After excluding non-chromosome-level sequences, as well as Y chromosomes, the number of BUSCO genes per-assembly ranged from 1820 in *Strongyloides ratti* to 3103 in *Caenorhabditis briggsae*, with only 961 being shared by all 14 genomes. We included all BUSCOs genes in the analysis, regardless of missingness. We next inferred rearrangements and ALGs across the phylogeny using the FF algorithm. The number of ALGs at each internal node varied between four and seven and we inferred a total of 16 fusions and 30 fissions. In contrast to Gonzalez de la Rosa *et al*. (2021), our analysis suggests the existence of six ALGs at the deepest node in the tree (the most recent common ancestor of Rhabditina and Tylenchina). These ALGs correspond to Nigon elements A, B, C, D and X with E + N fused (Figure 6). Another key difference between our results and those of Gonzalez de la Rosa *et al*. (2021) is that we infer a single fusion event of Nigons N + X in the ancestor of *Caenorhabditis sp*. and *Haemonchus contortus* (Figure 7), whereas they suggest that these ALGs fused independently in each of these lineages.

**Figure 6.**
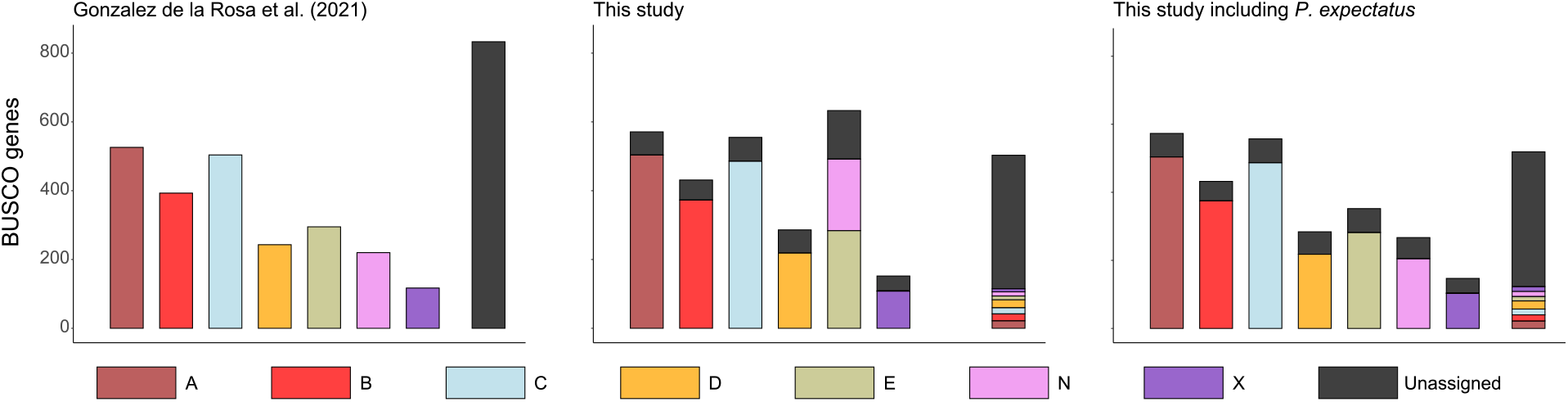
Ancestral linkage groups at the most recent common ancestor of Rhabditina and Tylenchina nematodes. Each plot shows ALGs as stacks of BUSCO genes, with the furthest right stack consisting of genes that were not assigned to any ALG. Stacks are coloured by the ALGs (i.e. Nigon elements) defined in Gonzalez de la Rosa *et al*. (2021) (left and legend below). Reconstructed ALGs in this study (middle) are similar to those of Gonzalez de la Rosa *et al*. (2021) but include a fusion of Nigons E + N. This fusion is not present, however, when including the genome of *Pristionchus exspectatus* in the analysis (right). Our method assigns more markers to ALGs than the clustering method of Gonzalez de la Rosa *et al*. (2021).

**Figure 7.**
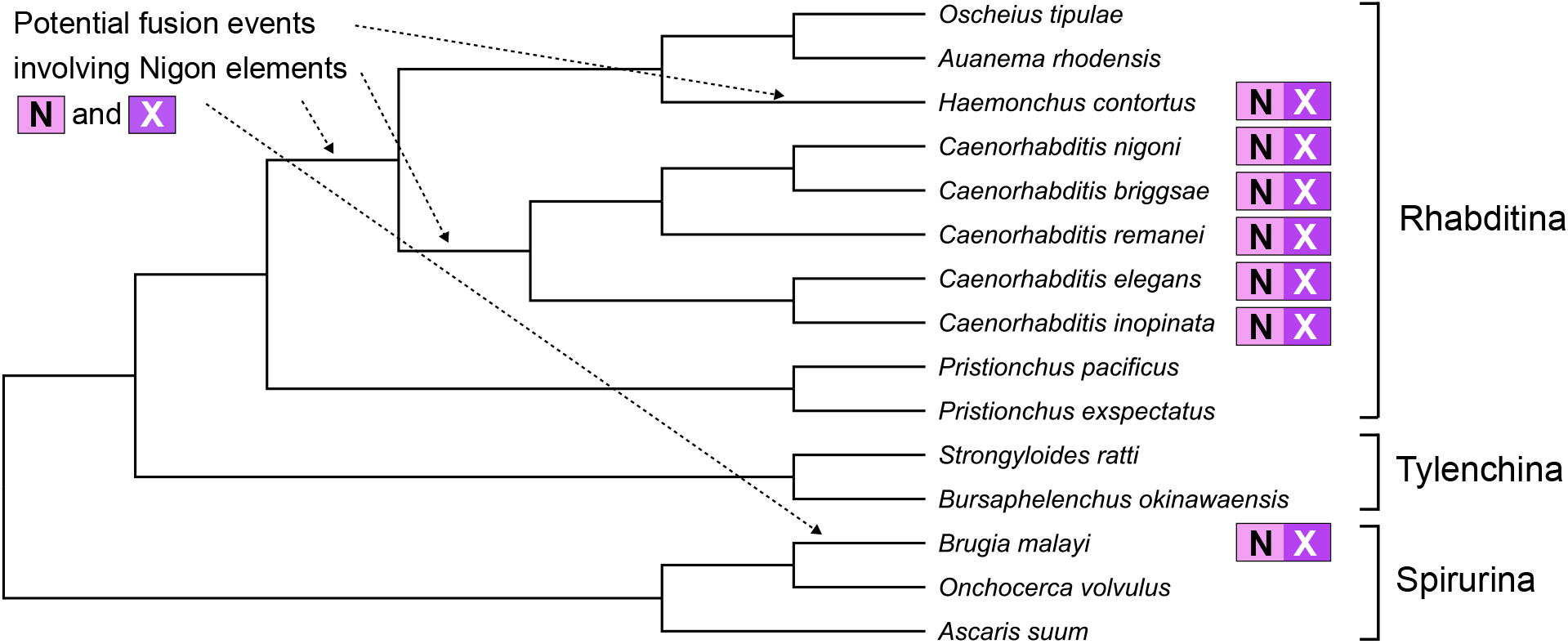
A tree showing the phylogenetic relationships between the 14 rhabditid nematode species analysed by Gonzalez de la Rosa *et al*. (2021), as well as *P. exspectatus*. The topology is from Gonzalez de la Rosa *et al*. (2021) and the branch lengths are arbitrary. Species with genomes where Nigon elements N + X are fused together are marked with a symbol to the right of the tree. Arrows point to branches where Nigon elements N + X could have fused.

The differences mentioned above can be explained by two limitations of our method. The first is that ancestral genomes are reconstructed locally, using information from only three other genomes at a time rather than the entire dataset. The reconstruction of Nigons E + N as one ALG deep in the tree, for example, can be explained by the fact that these elements co-occur on chromosomes of *Pristionchus pacificus* and *Strongyloides ratti*. Local reconstructions that rely on these genomes will therefore place Nigons E + N together, even though this necessitates multiple fission events on other branches of the phylogeny. The second limitation is that gene-order information is not used. The single fusion of Nigons N + X is supported by synteny but an examination of gene-order (as was done in Gonzalez de la Rosa *et al*. 2021) shows that these elements are well ‘mixed’ in *Caenorhabditis sp*. but much less so in *H. contortus* (see Figure 5 of Gonzalez de la Rosa *et al*. 2021). This is consistent with two independent fusion events.

Recently, Yoshida *et al*. (2023) investigated the evolution and consequences of chromosome fusions in *P. pacificus* and another closely related species in the genus, *P. exspectatus*. A comparison of these genomes shows that the E + N fusion in *P. pacificus* must be recent and not an ancestral state in rhabditid nematodes. Indeed, when we include the *P. exspectatus* genome assembly in our analysis we recover seven Nigon elements (A, B, C, D, E, N and X, Figure 6). So while limitations in our method can lead to incorrect inference, a denser sampling of present-day genomes does, unsurprisingly, improves performance.

Previous analyses of chromosome evolution in nematodes did not test for the presence of translocation rearrangements (Tandonnet *et al*. 2019; Gonzalez de la Rosa *et al*. 2021). To investigate this possibility, we performed inference allowing for translocation rearrangements while again including *P. exspectatus*. We inferred 42 rearrangements using the FFT algorithm (13 fusions, 24 fissions and 5 translocations), which is fewer than the 47 inferred with the FF algorithm (18 fusions and 29 fissions). We tested whether the five translocations inferred, representing 11.9% of the total rearrangements, could be a result of incorrect inference of a rearrangement history that only includes fissions and fusions. We simulated 18 fusions and 29 fissions over the phylogenetic tree (Figure 7) and inferred rearrangements using the FFT algorithm (see Methods for details). We find that only 2.7% of rearrangements are (incorrectly) inferred as translocations, with 95% CIs of 0.0 -10.3%. This provides some support for the idea that these nematode genomes have rearranged through translocation as well as fission and fusion (but see Discussion).

## Discussion

### Rearrangement rates and marker error

The need for efficient and accurate methods that infer past genome evolution will only increase as larger genome sequence datasets become available. Here we have implemented and evaluated one such class of method which focuses exclusively on synteny information. Our simulations show that it is straightforward to infer ALGs and rearrangements when the frequency of rearrangements per-branch is low relative to chromosome number. Although this will not always be the case, it is at least true in some groups of organisms. Most lepidoptera, for example, have a large number of chromosomes (*∼* 30) and slow rates of rearrangement, making inference straightforward (Wright *et al*. 2023). Higher rearrangement rates, however, impose a limit on the accuracy of reconstructed ALGs and rearrangements (Figures 2 and 3). The loss of synteny information with increasing rearrangement rates is hard to overcome within a parsimony-based framework, meaning that a different approach to modelling genome evolution may be required; e.g. estimating rearrangement parameters through simulation (Moshe *et al*. 2022) or performing a Bayesian sampling of rearrangement histories (Miklé s and Tannier 2010). Interestingly, Markov models of chromosome number evolution have recently been developed (Yoshida and Kitano 2021; Setter 2023) and could be adapted to estimate model parameters and sample likely rearrangement histories given a set of genomes. Alternative methods aside, our results show that high rearrangement rates hinder accurate inference and so we encourage researchers to consider the inherent uncertainty and limits to reconstructing rearrangement histories when genomes rearrange frequently.

We also explored the effect of marker errors on performance, finding that our method is robust to missing markers but not orthology error (Figure 4). Implementing a minimum set size did alleviate some of this effect, but this must be balanced against the risk of masking real rearrangements. We simulated orthology errors by moving markers between chromosomes, which should emulate incorrect orthology assignment as well as small scaffolding mistakes. It is unclear exactly how frequent these errors are in present-day genome assemblies and annotations. Anecdotally, initial assemblies of large genomes do often require extensive manual curation (e.g. Streicher *et al*. 2021), suggesting the possibility for small and overlooked scaffolding errors. Similarly, approximate methods for identifying orthology relationships, such a reciprocal best hits, can lead to false single copy orthologues (Emms and Kelly 2019). Limiting analyses to curated genome assemblies with high confidence single copy markers is therefore a sensible precaution when inferring ALGs and rearrangements.

### Lessons from a reanalysis of nematode genomes

Our analysis of nematode genomes highlighted some important limitations of our synteny-based inference method, `syngraph`. Firstly, `syngraph` only analyses a single triplet of genomes at a time, thereby ignoring useful synteny information contained in other genomes. The fact that we infer Nigons E + N as syntenic deep in the phylogeny (Figure 6) is a result of this limitation, as consideration of more genomes would generate a more parsimonious history involving two independent fusions of E + N on the lineages leading to *P. pacificus* and *S. ratti*. This problem could be alleviated by considering four or five closely related genomes at a time (at the expense of computation time) or by using a weighted graph as in Kim *et al*. (2017) and Muffato *et al*. (2023). An even simpler approach would be to perform iterative traversals of the tree until there is no further improvement in parsimony, although this would not guarantee a globally optimal solution (Adam and Sankoff 2008).

The second limitation highlighted by our analysis of nematode genomes is that we ignore useful information in the form of how mixed ALGs are in present-day genomes. Following a fusion of two ALGs, markers belonging to either ALG can become mixed through intra-chromosomal rearrangements. This mixing makes a fusion non-reversible as a subsequent fission is no longer likely to recover the original ALGs (Simakov *et al*. 2022; Schultz *et al*. 2023). Additionally, under the assumption that intra-chromosomal rearrangement rates are similar across a given clade, the amount of mixing between fused ALGs provides information about the timing of the fusion (Gonzalez de la Rosa *et al*. 2021). Without considering ALG mixing, it is difficult to infer whether Nigons N + X fused once in Rhabditina (and later fissioned apart in the lineage leading to *Oscheius tipulae* and *Auanema rhodensis*) or if they instead underwent two independent fusions (Figure 7). While our analysis with `syngraph` suggested the former, we agree with the interpretation of Gonzalez de la Rosa *et al*. (2021) that this fusion likely happened twice given patterns of ALG mixing in *Caenorhabditis sp*. and *H. contortus* and the improbability of a recent fission recovering two separate ALGs. This example suggests that there are limitations to considering synteny information alone and that including at least some gene-order information is likely to improve the accuracy of results.

Finally, we applied a simulation-based test for the presence of translocation rearrangements in the nematode dataset. The observed number of inferred translocations was greater than in 95% of simulations. While it is tempting to view this as strong evidence that the rearrangement history of nematode chromosomes involved translocations, it is important to remember that this result could be generated by other differences between the simulations and the rearrangement process underlying the real data. In particular, a non-uniform rate of rearrangement across the phylogeny (not captured by our simulations) could generate such a result, with (incorrectly) inferred translocations concentrated on branches of the tree with the most fissions and fusions. We therefore interpret this result as weak evidence for translocations and acknowledge the limitations of using simulations under highly simplified models that may only partially capture the complexities of real rearrangement histories.

### Gene tree discordance

We have so far assumed that the genealogies underlying rearrangements always follow the species tree. However, this assumption may not hold for groups of closely related populations / species due to appreciable levels of incomplete lineage sorting (ILS) and gene flow. Given that multiple chromosome-level assemblies are now being routinely generated for single species or genera (Kim *et al*. 2022; Liao *et al*. 2023; Shi *et al*. 2023), it is worth considering how gene tree discordance might affect our ability to accurately infer rearrangement histories. A rearrangement that has a history incongruent with the species tree will result in non-sister species carrying the derived chromosome arrangement (Jay *et al*. 2018). Assuming the species tree, this can be interpreted as two independent rearrangements or a single rearrangement with a subsequent reversion / loss. The challenge is therefore to discern between those scenarios as well the possibility of introgression / ILS. Useful evidence includes the probability of gene tree discordance estimated from polymorphism data (Dutheil *et al*. 2009), as well as whether rearrangement break points are consistent with multiple origins (Lundberg *et al*. 2023). A probabilistic method for rearrangement inference that includes gene tree discordance seems like a distant goal at present, but will be useful in any analysis where rearrangements happen on the same time scale as lineage sorting.

### Outlook

We have investigated how synteny information can be used to estimate past genome evolution, co-estimating both ALGs and rearrangements. Genome rearrangement problems have garnered considerable interest within the field of mathematics (Fertin *et al*. 2009), but here we have focused on a relatively simple version of the problem as well as the practicalities of analysing real data. We have been motivated by the fact that accurate inference of past rearrangements has the ability to improve our understanding of how genomes evolve. For example, we still do not know the relative fitness effects of different types of rearrangements (e.g. sex-autosome fusions vs. autosome-autosome fusions) or whether the majority of new fission and fusions are weakly deleterious (Pennell *et al*. 2015). Additionally, while the importance of fissions and fusions in speciation is becoming clearer (Yoshida *et al*. 2023; Mackintosh *et al*. 2023), we still have an incomplete understanding of exactly how such rearrangements prevent gene flow. Identifying inter-chromosomal rearrangements across species and populations will be the first step in answering these biological questions, and it is encouraging that our synteny-based method has already been used to generate new results about genome evolution (Mackintosh *et al*. 2023; Wright *et al*. 2023). We anticipate that a variety of methods, focusing on different rearrangements types and relying on different inference procedures, will be required to investigate genome evolution across the tree of life and improve our understanding of the role of chromosome rearrangements in evolution.

## Methods

### An overview of syngraph

We implemented methods for investigating past genome evolution from synteny in a modular python tool, `syngraph` (https://github.com/A-J-F-Mackintosh/syngraph). The suggested workflow is to first generate an adjacency graph from orthology data using the `syngraph build` module. A file containing markers and their chromosome assignments must be provided for each genome, with matching marker IDs denoting orthology across genomes. Given the adjacency graph and a phylogenetic tree, ALGs and rearrangements can then be estimated with `syngraph infer`. This generates descriptions of rearrangements across the phylogeny as well as a new graph which includes reconstructed ALGs. This graph can be summarised with `syngraph tabulate`, which produces a table with the assignment of each marker to an ALG.

### ALG reconstruction under missingness and error

For a triplet of genomes, *G*_*A*_, *G*_*B*_, *G*_*C*_, the ALGs at *G*_*i*_ (the genome at the internal node of the tree connecting them) are estimated by considering synteny-sets. An adjacency-based approach builds ALGs by combining synteny-sets that are syntenic in *>* 2 genomes, whereas an event-based finds the set of ALGs that minimises the syntenic distance between *G*_*i*_ and *G*_*A*_, *G*_*B*_, *G*_*C*_. However, ALG reconstruction must be modified when some markers are missing. If a marker is missing from *G*_*C*_ then the synteny-set for such a marker would only have a partial notation, e.g. [*A*_4_, *B*_22_] rather than [*A*_4_, *B*_22_, *C*_9_]. One solution is to limit the analysis to markers shared across all genomes, with full notations, but this quickly becomes restrictive as the number of leaves in the phylogeny grows. We instead attempt to assign markers with partial missingness to ALGs. Specifically, a marker with notation [*A*_4_, *B*_22_] can be assigned if there is a single (already reconstructed) ALG with notation [*A*_4_, *B*_22_, *C*_*∗*_], with *∗* representing any sequence in *G*_*C*_. Markers that are missing from two genomes, or those that are not consistent with a single ALG, are not assigned. This procedure is performed by default within `syngraph infer`, but can be disable by restricting the analysis to markers present in all genomes when reading in data with `syngraph build`.

We also considered the effect of orthology error and how ALG estimation can be made more robust to it. Synteny-sets will vary in size depending on: (i) the total number of markers in the analysis, (ii) the size of chromosomes, (iii) the number of rearrangements between genomes, and (iv) the frequency of orthology error. As an example, consider a chromosome that is conserved in a triplet of genomes, so that hundreds of markers have the same notation, e.g. [*A*_4_, *B*_22_, *C*_9_], forming a large synteny-set. If a gene on sequence *C*_15_ is falsely annotated as having orthology with genes on sequences *A*_4_ and *B*_22_ we now obtain a new synteny-set with notation [*A*_4_, *B*_22_, *C*_15_]. Importantly, this synteny set is small, consisting of only a single marker. We therefore implemented an option in `syngraph infer` to enforce a minimum size of synteny-sets (`--minimum`), with the aim of minimising the effect of orthology / scaffolding error.

### Simulations

To investigate the performance of these methods we simulated rearrangement histories and attempted to infer them with `syngraph`. The general simulation procedure is as follows: For each simulation a phylogenetic tree is generated under a birth-death process using Dendropy (Sukumaran and Holder 2010). The birth rate is set to one 1.0 and the death rate is 0.5. The tree is sampled once there are *n* + 1 leaves. As a result, two leaves will have external branch lengths of zero and so one of them is removed to recover a tree with *n* leaves. This means that the time of sampling is effectively a random event determined by the time it takes for the number of leaves to increase from *n* to *n* + 1. A genome with *k* chromosomes is initiated at the root of the tree, with *g* markers uniformly distributed among them. A total of *r* rearrangements are placed onto branches of the tree, with probability proportional to branch lengths and different rearrangement types being sampled under a specific fission:fusion:translocation ratio. The initial genome at the root is then simulated forwards in time across the tree and rearranged through set operations. The resulting markers at the leaves of the tree and the phylogeny are parsed to `syngraph infer`. Unless stated otherwise, simulations were parameterised with *k* = 20 initial chromosomes and *g* = 1000 markers. Simulations involving only fission and fusions were simulated with rearrangement ratio 1:1:0, whereas those involving fissions, fusions, and translocations were simulated with ratio 1:1:1 (unless otherwise stated). Performance metrics were always estimated using 1,000 simulations for a given parameter combination.

Marker missingness was introduced by randomly removing *m*% of markers per-genome before parsing the markers to `syngraph infer`. Similarly, marker orthology error was introduced by randomly selecting *e*% of markers per-genome, and then placing each selected marker on a chromosome with uniform probability. This allowed for the possibility of a selected marker being placed on the chromosome from which it was sampled.

### Reanalysis of nematode genomes

We reanalysed 14 rhabditid nematode genomes (Table S1) and also performed analyses including the genome sequence of *Pristionchus exspectatus*. For each genome assembly we identified sequences corresponding to nematoda odb10 single-copy genes using BUSCO v5.2.2 (Manni *et al*. 2021).

BUSCO genes annotated on non-chromosome-level sequences or Y chromosomes were removed from the analysis. We then performed the `syngraph` workflow described above using a phylogenetic tree with topology from Gonzalez de la Rosa *et al*. (2021) and branch lengths corresponding to at least one time unit between speciation events. Missingness was allowed when reading in markers and the minimum synteny-set size for inference was set to 20.

We also performed a simulation test for translocation rearrangements. Each simulation (1,000 in total) had an initial genome with 7 chromosomes, 1,000 markers, and a total of 47 rearrangements with fission:fusion:translocation ratio 29:18:0. The simulations were conditioned on the phylogenetic tree presented in Figure 7, and the minimum synteny-set size was again set to 20. The proportion of inferred rearrangements that were translocations were recorded for each simulation. The mean was calculated across simulations and the 95% CIs were estimated using the 25th and 975th percentiles.

## Acknowledgments

We would like to thanks Derek Setter and Charlotte Wright for comments on an earlier version of this manuscript. AM is supported by an E4 PhD studentship from the Natural Environment Research Council (NE/S007407/1). KL was supported by a fellowship from the Natural Environment Research Council (NERC, NE/L011522/1) and a European Research Council starting grant (ModelGenomLand 757648). SHM is supported by a Royal Society University Research Fellowship (URF/R1/180682).

## Data accessibility

The command-line tool, `syngraph`, is available at https://github.com/A-J-F-Mackintosh/syngraph. Scripts for simulating rearrangement histories are available at the same Github directory. The NCBI accessions for genome assemblies analysed in this work are given in Table S1.

