## Supplementary material for "Inferring inter-chromosomal rearrangements and ancestral linkage groups from synteny"

### Supplementary Materials

#### Supplementary Methods

The heuristic algorithm of Ferretti *et al.* (1996) finds the syntenic distance between two genomes using fission, fusion and translocation rearrangements. Their algorithm allows non-reciprocal translocations, which are unidentifiable from a fission and subsequent fusion. In this work we only consider reciprocal translocations, as they are more likely to be identifiable from fissions and fusions (Figure 5). We therefore modified the algorithm of Ferretti *et al.* (1996) to only include reciprocal translocations. Here we describe the algorithm through a simple example.

Consider two genomes, $G_A$ and $G_B$, that consist of chromosome containing orthologous markers (lower case letters). We can write these genomes as

$G_A$: [a b c d e f] [g h i] [j k] [l m n o p q r]

$G_B$: [a b c q r] [g h i j k] [d e f l m] [n o p]

Using the compact representation of Ferretti *et al.* (1996), we can instead write the chromosomes of $G_A$ in terms of the $G_B$ chromosomes that they share markers with.

$G_A$: $[B_1\ B_3]$ $[B_2]$ $[B_2]$ $[B_3\ B_4\ B_1]$ [a b c d e f] [g h i] [j k] [l m n o p q r]

To transform $G_A$ into $G_B$, we first identify labels (highlighted in bold below) that are only present once and implement a fission to generate a new chromosome containing that label.

$[B_1\ B_3]$ $[B_2]$ $[B_2]$ $[B_3\ \mathbf{B_4}\ B_1]$ [a b c d e f] [g h i] [j k] [l m n o p q r]

↓ fission ↓

$[B_1\ B_3]$ $[B_2]$ $[B_2]$ $[B_3\ B_1]$ $[\mathbf{B_4}]$ [a b c d e f] [g h i] [j k] [l m q r] [n o p]

If there are no more labels present only once, we choose a label that is present twice. If the chromosomes that share that label also share at least one other label, then a translocation is implemented.

$$[\mathbf{B_1}\ B_3] \quad [B_2] \quad [B_2] \quad [B_3\ \mathbf{B_1}] \quad [B_4] \qquad [\text{a b c d e f}] \quad [\text{g h i}] \quad [\text{j k}] \quad [\text{l m q r}] \quad [\text{n o p}]$$

$$\downarrow \text{translocation} \downarrow$$

$$[\mathbf{B_1}] \quad [B_2] \quad [B_2] \quad [B_3] \quad [B_4] \qquad [\text{a b c q r}] \quad [\text{g h i}] \quad [\text{j k}] \quad [\text{d e f l m}] \quad [\text{n o p}]$$

If we have chosen another label that is present twice, but the chromosomes that share that label do not share any other labels, then a fusion is implemented.

$$[B_1] \quad [\mathbf{B_2}] \quad [\mathbf{B_2}] \quad [B_3] \quad [B_4] \qquad [\text{a b c q r}] \quad [\text{g h i}] \quad [\text{j k}] \quad [\text{d e f l m}] \quad [\text{n o p}]$$

$$\downarrow \text{fusion} \downarrow$$

$$[B_1] \quad [\mathbf{B_2}] \quad [B_3] \quad [B_4] \qquad [\text{a b c q r}] \quad [\text{g h i j k}] \quad [\text{d e f l m}] \quad [\text{n o p}]$$

We have now recovered $G_B$ through rearrangement and so there are no more steps. The syntenic distance is three, and the putative rearrangements history involves one fission, one translocation and one fusion.

In the above example we did not encounter any instances where there were labels shared by $> 2$ chromosomes. In this case, a fusion is implemented involving the two chromosomes with the greatest intersection of labels.

Table S1: Nematode genomes analysed in this study. The first 14 taxa listed were originally analysed by Gonzalez de la Rosa *et al.* (2021), whereas the *Pristionchus exspectatus* genome was not. In some cases more recent assemblies were used than in Gonzalez de la Rosa *et al.* (2021).

| **Taxon** | **Chr.** | **NCBI accession** | **Study** |
|---|---|---|---|
| *Ascaris suum* | 24 | GCA_013433145.1 | (Wang *et al.* 2020) |
| *Auanema rhodensis* | 7 | GCA_947366455.1 | (Tandonnet *et al.* 2019) |
| *Brugia malayi* | 5 | GCA_000002995.5 | (Tracey *et al.* 2020) |
| *Bursaphelenchus okinawaensis* | 6 | GCA_904067145.1 | (Sun *et al.* 2020) |
| *Caenorhabditis briggsae* | 6 | GCA_021491975.1 | (Stevens *et al.* 2022) |
| *Caenorhabditis elegans* | 6 | GCA_028201515.1 | (Bush *et al.* 2023) |
| *Caenorhabditis inopinata* | 6 | GCA_003052745.1 | (Kanzaki *et al.* 2018) |
| *Caenorhabditis nigoni* | 6 | GCA_027920645.1 | NA |
| *Caenorhabditis remanei* | 6 | GCA_010183535.1 | (Teterina *et al.* 2020) |
| *Haemonchus contortus* | 6 | GCA_000469685.2 | (Doyle *et al.* 2020) |
| *Onchocerca volvulus* | 4 | GCA_000499405.2 | (Cotton *et al.* 2016) |
| *Oscheius tipulae* | 6 | GCA_013425905.1 | (Gonzalez de la Rosa *et al.* 2021) |
| *Pristionchus pacificus* | 6 | GCA_000180635.4 | (Rödelsperger *et al.* 2017) |
| *Strongyloides ratti* | 3 | GCA_001040885.1 | (Nemetschke *et al.* 2010) |
| *Pristionchus exspectatus* | 6 | GCA_911812115.1 | (Yoshida *et al.* 2023) |
